# Phospho-ERK is a response biomarker to a combination of sorafenib and MEK inhibition in liver cancer

**DOI:** 10.1101/252452

**Authors:** Cun Wang, Haojie Jin, Dongmei Gao, Bastiaan Evers, Guangzhi Jin, Zheng Xue, Liqin Wang, Roderick Beijersbergen, Wenxin Qin, René Bernards

**Affiliations:** Division of Molecular Carcinogenesis, Oncode Institute. The Netherlands Cancer Institute, Plesmanlaan 121, 1066 CX Amsterdam, The Netherlands.; State Key Laboratory of Oncogenes and Related Genes, Shanghai Cancer Institute, Renji Hospital, Shanghai Jiao Tong University School of Medicine, Shanghai, China; Liver Cancer Institute, Zhongshan Hospital, Fudan University, Key Laboratory of Carcinogenesis and Cancer Invasion, Ministry of Education, Shanghai, China.; Department of Pathology, Eastern Hepatobiliary Surgery Hospital, Second Military Medical University, Shanghai, China.

**Keywords:** Hepatocellular carcinoma, CRISPR screen, sorafenib, MAPK1

## Abstract

Treatment of liver cancer remains challenging, due to a paucity of drugs that target critical dependencies. Sorafenib is a multikinase inhibitor that is approved as the standard therapy for advanced hepatocellular carcinoma patients, but it can only provide limited survival benefit for patients. To investigate the cause of this limited therapeutic effect, we performed a CRISPR-Cas9 based synthetic lethality screen to search for kinases whose knockout synergize with sorafenib. We find that suppression of ERK2 sensitizes several liver cancer cell lines to sorafenib. Drugs inhibiting the MEK or ERK kinases reverse unresponsiveness to sorafenib *in vitro* and *in vivo* in a subset of liver cancer cell lines characterized by high levels of active phospho-ERK levels through synergistic inhibition of ERK kinase activity. Our data provide a combination strategy for treating liver cancer and suggest that tumors with activation of p-ERK, which is seen in some 30% of liver cancers, are most likely to benefit from such combinatorial treatment.

## Introduction

Liver cancer is one of the most frequent malignancies and the second leading cause of cancer-related deaths worldwide (Llovet et al, 2016). In the last decade, our understanding of the genetic landscape of hepatocellular carcinoma (HCC) has improved significantly through large-scale sequencing studies. As a result, we now know that several signaling pathways are involved in HCC initiation and progression, including telomere maintenance, WNT-β-catenin pathway, cell cycle regulators, epigenetic regulators, AKT/mTOR and MAPK pathway (Zucman-Rossi et al, 2015). However, the most frequent mutations in HCC are currently undruggable. Sorafenib is a multi-kinase inhibitor that is approved as the standard therapy for advanced HCC patients, but only provides 2.8 months survival benefit for patients with advanced HCC (Llovet et al, 2008). The survival advantage is even more limited for Asia-Pacific patients (2.3 months) (Cheng et al, 2009). Regorafenib is a treatment strategy shown to provide overall survival benefit in HCC patients progressing on sorafenib treatment (Bruix et al, 2017). However, chemically, regorafenib and sorafenib differ by just one atom (Gyawali & Prasad, 2018). The regorafenib registration trial design involved a high degree of patient selection. Even in such ideal settings, only modest clinical benefit was achieved. Lenvatinib is also a multi-kinase inhibitor, which provided impressive antitumor activity in phase II trials in patients with advanced unresectable HCC (Ikeda et al, 2017). However, there is only 1.3 months additional survival benefit for patients compared to sorafenib in the phase III trials. These clinical data point to the need to reconsider the current therapeutic strategy. A recurrent problem in clinical studies in liver cancer is the paucity biomarkers linked to drug response (Gerbes et al, 2017).

Combination therapies can help fight therapy failure and improve response to approved drugs. We have successfully employed functional genetic screens to find powerful combinations of cancer drugs by exploiting the concept of ‘synthetic lethality’. This has resulted in clinical testing of several combination treatment strategies, including the use of BRAF inhibitor combined with EGFR inhibitor in *BRAF(V600E)* mutant colon cancer (Prahallad et al, 2012) (NCT 01719380, NCT01791309, NCT02928224) and MEK inhibitor combined with EGFR/ERBB2 inhibitor in *KRAS* mutant cancer (Sun et al, 2014) (NCT 02039336, NCT02230553: NCT02450656). Some of these studies have already revealed promising clinical activity in these difficult-to-treat patient populations (van Geel et al, 2017). For liver cancer, the concept of ‘synthetic lethality’ has also been validated by the combination of sorafenib and MAPK14 inhibition, which was identified by an *in vivo* RNAi screen (Rudalska et al, 2014).

CRISPR-Cas9 is a powerful gene-editing technology that has been widely employed for genome-scale functional screens (Sanchez-Rivera & Jacks, 2015). Compared to the traditional shRNA-based system, CRISPR technology provides advantages in low noise, minimal off-target effects and high consistent activity (Evers et al, 2016). Taking advantage of the CRISPR-Cas9 based functional screening system, we developed a platform that can be used to identify kinases whose inhibition increases the therapeutic efficacy of cancer drugs.

We describe here the use of kinome-centered synthetic lethality screens to identify enhancers of the response to sorafenib in liver cancer. Moreover, we identify a biomarker of response to the drug combination identified here, which may be useful in the future clinical development of this drug combination.

## Results and Discussion

### HCC cell lines are unresponsive to sorafenib

To study how liver cancer cells respond to sorafenib *in vitro*, we determined the efficacy of sorafenib in nine liver cancer cell lines using a long-term proliferation assay. Fig 1A shows all cell lines are relatively insensitive to sorafenib, in agreement with the modest clinical benefit of this drug (Cheng et al, 2009; Llovet et al, 2008). Sorafenib is approved as a treatment for advanced renal cell carcinoma, unresectable HCC and thyroid cancer. We therefore further analyzed the efficacy of sorafenib in cell lines from these different cancer types in the cancer cell line encyclopedia (CCLE) (Barretina et al, 2012). Consistent with our results, the vast majority of these cancer cell lines have an IC50 for sorafenib of over 5 μM (Appendix Fig S1A). The highly related drug regorafenib was approved by the FDA as a second-line therapy for locally advanced or metastatic HCC patients with disease progression after sorafenib treatment (Bruix et al, 2017). In a long-term proliferation assay, we found that the vast majority of HCC cell lines are also insensitive to regorafenib (Appendix Fig S1B). Together, both the cell line data and the clinical data indicate a modest activity of sorafenib or regorafenib in liver cancer.

**Figure 1.**
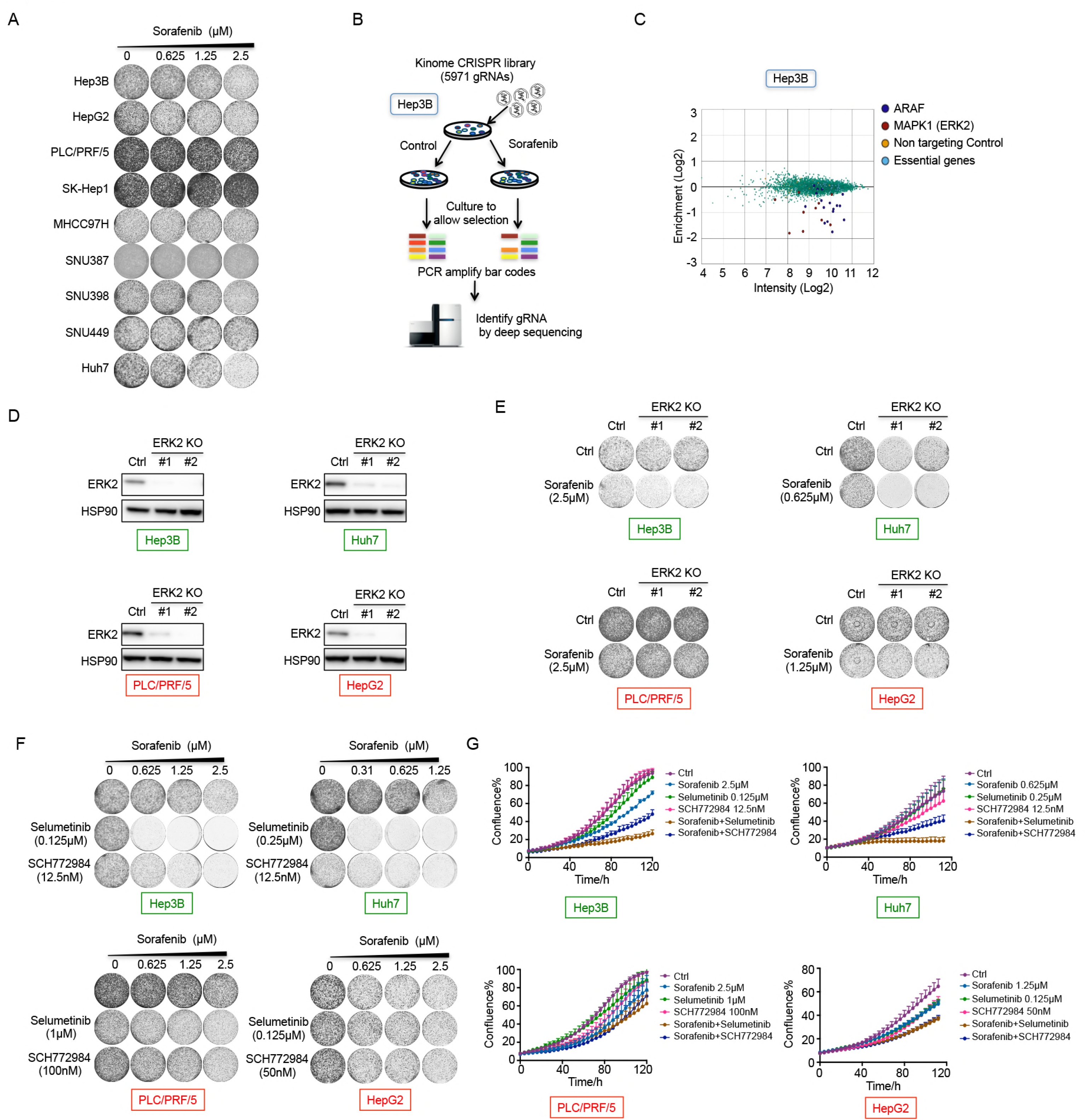
A synthetic lethal screen identifies that ERK2 inhibition confers sensitivity to sorafenib in liver cancer cell lines. A Liver cancer cell lines were treated with increasing concentrations of sorafenib for about two weeks. Viability was assessed by a colony formation assay. B Schematic outline of the synthetic lethal screen. Pooled kinome CRISPR screen were performed in Hep3B cells. C Representation of the relative abundance of the gRNA barcode sequences from the synthetic lethal screen. The y axis shows enrichment (relative sorafenib treated / untreated) and x axis shows intensity (average sequence reads in untreated sample) of each gRNA. *ARAF* and *MAPK1* were identified as synthetic lethal hits in Hep3B cells. D The level of ERK2 knockout was determined by western blot. HSP90 protein level served as a loading control. E Two independent gRNAs targeting *ERK2* enhance response to sorafenib in Hep3B and Huh7 cells, but not in PLC/PRF/5 and HepG2 cells. These cells were culture with or without sorafenib for 10 days. F-G Long-term colony formation and short-term IncuCyte^®^ cell proliferation assay show synergistic response of sorafenib combined with MEK or ERK inhibitor in Hep3B and Huh7 cells, but not in PLC/PRF/5 and HepG2 cells.

### A synthetic lethal screen with sorafenib

We have described the use of kinome-centered RNA interference (RNAi) genetic screens, which enable the identification of kinases whose inhibition is synergistic with the BRAF inhibitor in *BRAF* mutant colon cancer and MEK inhibition in *KRAS* mutant lung and colon cancer (Prahallad et al, 2012; Sun et al, 2014). To systematically identify the kinases whose inhibition confers sensitivity to sorafenib in liver cancer cells, Hep3B cells were infected with the lentiviral kinome gRNA collection and cultured in the absence or presence of sorafenib for 14 days. After this, genomic DNA was isolated from both treated and untreated cells, and the relative abundance of gRNA was determined by next generation sequencing of the bar code contained in each gRNA vector in three biological replicates (Fig 1B and Appendix Fig S2A). Several independent gRNA vectors targeting *ARAF* and *MAPK1* (*ERK2*) were identified from the synthetic lethal screen (Fig 1C and Appendix Fig S2B). To validate these screen hits, we first treated liver cancer cell lines (Hep3B, SNU398, HepG2, and PLC/PRF/5) with a combination of sorafenib and LY3009120 (a pan-RAF inhibitor). Synergy between these two drugs was seen only in the Hep3B cells used in the screen, but in none of three other HCC cell lines (Appendix Fig S3A). Similar results were seen when we treated liver cancer cell lines (Hep3B, SNU398, HepG2, and PLC/PRF/5) with a combination of regorafenib and LY3009120 (Appendix Fig S3B). We conclude that the synthetic lethal effects of *ARAF* knockout and sorafenib may be context-dependent and therefore unlikely to be of general utility in HCC treatment. Next, we focused on the ERK2 hit identified in the screen. We infected liver cancer lines (Hep3B, Huh7, PLC/PRF/5, and HepG2) with two *ERK2* gRNA vectors (both of them decreased ERK2 levels [Fig 1D]). These cells were then cultured with or without sorafenib for 10 days. Suppression of ERK2 in combination with sorafenib caused an obvious inhibition of proliferation in Hep3B and Huh7 cells, but not in PLC/PRF/5 and HepG2 cells (Fig 1E).

To study whether MAPK signaling could be responsible for the poor response to sorafenib and the synergy with ERK2 deficiency, we treated these four liver cancer cell lines with a combination of sorafenib and selumetinib (MEK inhibitor) or the combination of sorafenib and SCH772984 (ERK inhibitor). Similar to what was observed upon knockout of ERK2, selumetinib and SCH772984 each showed strong synergy with sorafenib in Hep3B and Huh7 cells (Fig 1F-G), but a synergistic effect was not observed in PLC/PRF/5 and HepG2 cells (Fig 1F-G). Similar results were seen for the combination of regorafenib and selumetinib (Appendix Fig S4A-B). These results suggest that cell-intrinsic mechanisms exist that determine the response to the combination of sorafenib and MEK inhibition.

### Synergistic inhibition of p-ERK causes apoptosis in HCC cell lines

To address the mechanism by which sorafenib and selumetinib synergize to reduce viability of several liver cancer cell lines, we assayed induction of apoptosis in the presence of sorafenib, selumetinib or the combination of the two drugs. Both Hep3B and Huh7 cells showed modest evidence of apoptosis following monotherapy. However, strong synergistic induction of apoptosis was observed in these cells when sorafenib and selumetinib were combined, as indicated by the IncuCyte^®^ caspase-3/7 apoptosis assay (Fig 2A). Consistent with modest effects on inhibition of proliferation in PLC/PRF/5 and HepG2 cells, sorafenib and selumetinib combination displayed no or little evidence on apoptosis induction. Moreover, biochemical analysis indicated that the drug combination resulted in synergistic inhibition of p-ERK in Hep3B and Huh7 cells. However, we cannot detect p-ERK in HepG2 and PLC/PRF/5 cells, even in the absence of any drug (Fig 2B). Similar results were observed when we treated liver cancer cell lines with the combination of regorafenib and selumetinib (Appendix Fig S4C-D).

**Figure 2.**
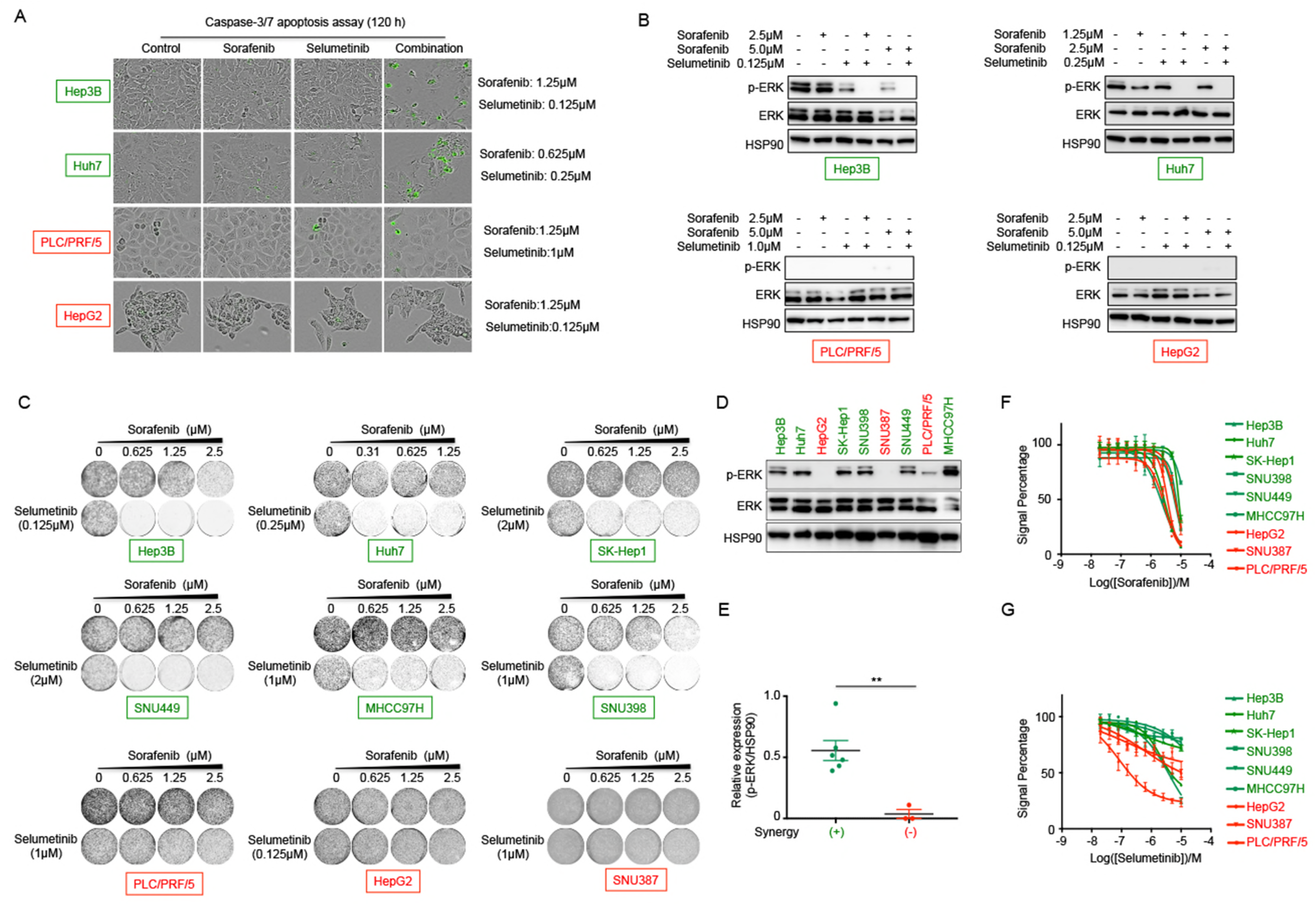
phospho-ERK determines synthetic lethal effects of sorafenib and MEK inhibition in liver cancer. A Representative images of sorafenib, selumetinib or the combination of both drugs treated cells in the presence of a caspase-3/7 activatable dye. B Biochemical responses of HCC cell lines treated with sorafenib, selumetinib or their combination were documented by western blot analysis. Cells were harvested at 48 h after drug treatment. C The response of liver cancer cell lines to the combination of sorafenib and selumetinib. D Western blot analysis of p-ERK and ERK levels in a panel of liver cancer cell lines. HSP90 protein level served as a loading control. E High p-ERK levels correlates with high synergistic response to the treatment containing sorafenib and selumetinib. \*\*\**P* < 0.001. F Short-term growth-inhibition assay of a panel of liver cancer cell lines to sorafenib treatment. G Short-term growth-inhibition assay of a panel of liver cancer cell lines to selumetinib treatment.

### Biomarker of response to the combination therapy

To ask whether p-ERK could be a potential biomarker for the drug combination treatment, we determined their synthetic lethal effects in 9 liver cancer cell lines. The combination of sorafenib and selumetinib induced strong synergistic effects in six cell lines having high p-ERK. However, at best modest effects were observed in the three cell lines having lower intrinsic p-ERK (HepG2, SNU387, and PLC/PRF/5) (Fig 2C-E). There are several studies indicating that baseline p-ERK may be a predictive biomarker of clinical response to sorafenib (Chen et al, 2013; Zhang et al, 2009). We note that in the panel of our HCC cell lines, there is not a strong correlation between the baseline p-ERK and sensitivity to sorafenib or selumetinib monotherapy (Fig 2F-G). Together, our data suggest a combination therapy for the treatment of liver cancer that is most likely to be effective in tumors having high basal p-ERK levels.

Several preclinical studies and two clinical trials have been performed in liver cancer with the combination of sorafenib and MEK inhibitor (selumetinib or refametinib) (Lim et al, 2014; Schmieder et al, 2013; Tai et al, 2016). Both combinations are tolerable by most patients with encouraging anti-tumor activity. The median overall survival was 14.4 months in the phase Ib study of selumetinib in combination with sorafenib in advanced HCC patients, but there was no biomarker identified predictive of response (Tai et al, 2016). In a previous study, activation of p-ERK was identified in only 10.3% of HCC (Newell et al, 2009). We also analyzed p-ERK using a tissue microarray (TMA) containing 78 HCC specimens by immunochemical analysis. Levels of p-ERK in tumor tissues were classified as high expression in 11 cases (11/78, 14.1%), low expression in 11 cases (11/78, 14.1%), or negative in 56 cases (56/78, 71.8%) (Fig 3A). Our finding suggests that patient selection based on presence of p-ERK (seen in about 30% patients) before initiating combination treatment may be useful to identify responders to this combination.

**Figure 3.**
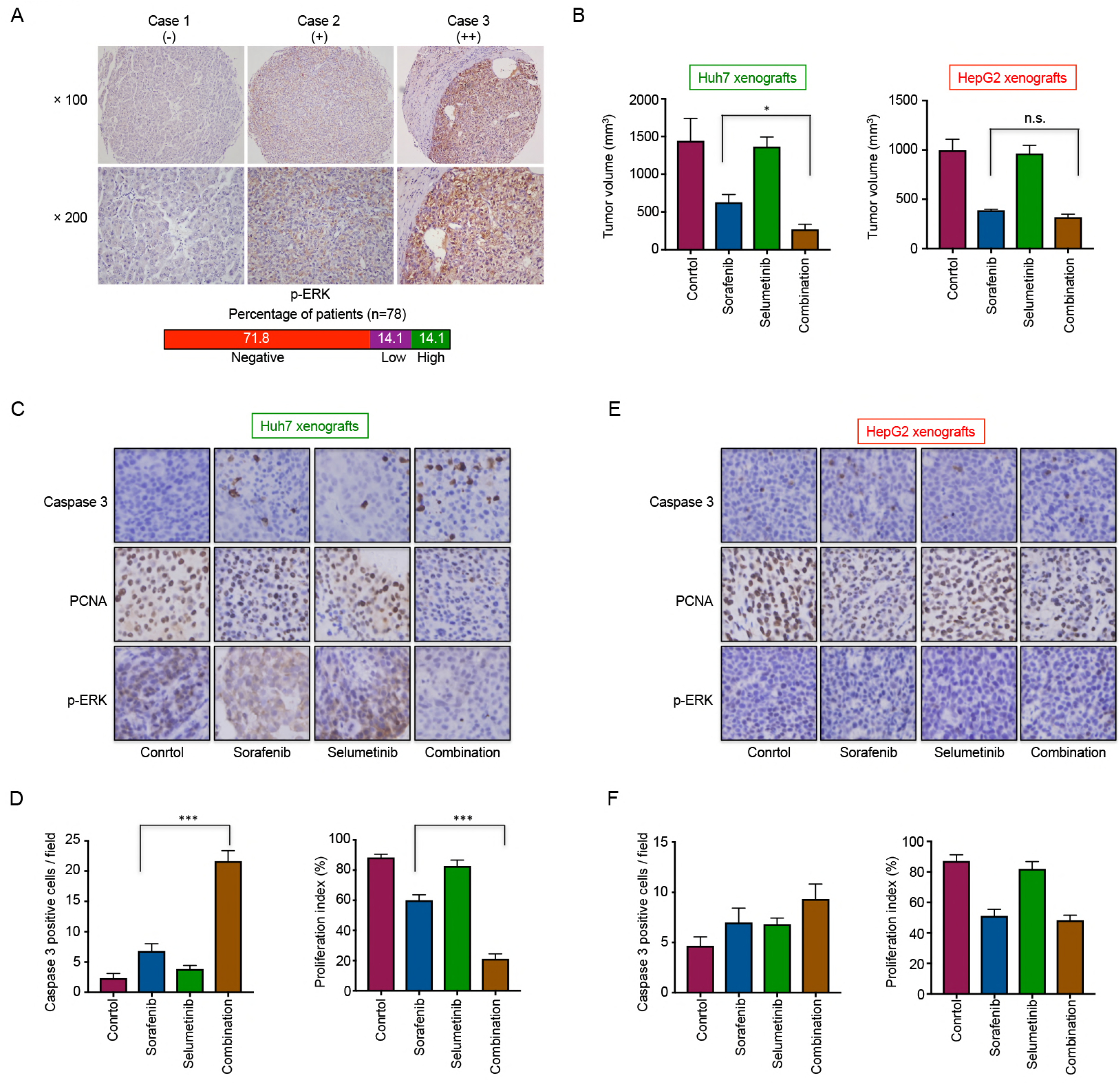
Sorafenib and selumetinib specifically synergize to suppresses *in vivo* growth of p-ERK activated xenograft models. A Typical images of p-ERK in HCC tissues by immunohistochemical staining (scored as high expression, low expression, or negative). B Combination of sorafenib and selumetinib can selectively synergize to suppress tumor growth in p-ERK activated xenograft models. \**P* < 0.05; n.s., not significant. C-D Immunohistochemical staining of caspase 3, p-ERK and PCNA in Huh7 xenografts (magnification, ×400). The number of caspase 3 positive cells and PCNA positive cells were quantified. \*\*\**P* < 0.001. E-F Immunohistochemical staining of caspase 3, p-ERK and PCNA in HepG2 xenografts (magnification, ×400). The number of caspase 3 positive cells and PCNA positive cells were quantified.

### *In vivo* effects of the combination therapy

To assess whether the *in vitro* findings can be recapitulated *in vivo*, Huh7 (high p-ERK level) and HepG2 (low p-ERK level) cells were injected into nude mice. Upon tumor establishment, xenografts were treated with vehicle, sorafenib, selumetinib, or the combination for about 21 days. As shown in Fig 3B, the combination of sorafenib and selumetinib elicited a potent growth inhibition of Huh7 cells. However, no synergistic effects were observed in HepG2 xenografts, which have no detectable p-ERK level *in vitro*.

To confirm the mechanism identified *in vitro*, the effect of sorafenib and selumetinib on p-ERK in Huh7 xenografts was assayed by immunohistochemical staining. Treatment of mice with sorafenib or selumetinib alone result in limited suppression of p-ERK. In contrast, the combination treatment showed a potent inhibition on activation of ERK accompanied with increase in caspase 3 positive cells and decrease in PCNA (a proliferation marker) positive cells (Fig 3C-D). Similar to the *in vitro* experiments, we could not detect p-ERK in HepG2 xenografts (Fig 3E-F).

Our data provide a strong rationale for the use of a combination of sorafenib and MEK inhibitor in liver cancer patients with high p-ERK level. The strong synergistic effect between sorafenib and MEK inhibitor described here is explained by a powerful suppression of activation of ERK. Moreover, p-ERK could be a potentially powerful predictive biomarker of response to the combination therapy. Whether baseline p-ERK can serve as a useful predictive biomarker for the response to the combination treatment needs to be addressed in future prospective clinical studies, as suitable tissue from existing clinical studies is not available for analysis.

## Materials and Methods

### Cell lines

The human HCC cell lines, Hep3B, HepG2, SNU387, SK-Hep1, SNU398, SNU449, and PLC/PRF/5 were purchased from the American Type Culture Collection (ATCC, VA, USA). Huh7 cells were purchased from Riken Cell Bank (Tsukuba, Japan). MHCC97H was provided by the Liver Cancer Institute of Zhongshan Hospital (Shanghai, China). HCC cells were cultured in DMEM with 10% FBS, glutamine and penicillin/streptomycin (Gibco^®^) at 37 °C / 5% CO_2_. Mycoplasma contamination was excluded via a PCR-based method. The identities of all the cell lines were confirmed by STR testing.

### Compound and antibodies

Sorafenib was purchased MedKoo Bioscience. Selumetinib, SCH772984, and LY3009120 were purchased from Selleck Chemicals. Antibody against HSP90 (H-114) and ERK2 (sc-154) were purchased from Santa Cruz Biotechnology. Antibodies against p-ERK (4695) and ERK (9101) were purchased from Cell Signaling Technology.

### Pooled ‘synthetic lethal’ CRISPR screen

For the design of the kinome CRISPR library, 5921 gRNAs targeting 504 human kinases, 10 essential genes, and 50 non-targeting gRNAs were selected (Appendix Table S1). Oligo’s with gRNA sequences flanked by adapters were ordered from CustomArray Inc (Bothell, WA) and cloned as a pool by GIBSON assembly in LentiCRISPRv2.1. The kinome CRISPR library was introduced to Hep3B cells by lentiviral transduction. Cells were then pooled and plated at 500,000 cells per 15 cm dish in the absence or presence of 1.5 μM sorafenib (12 dishes for each condition) and the medium was refreshed twice per week for 14 days. The abundance of each gRNA in the pooled samples was determined by Illumina deep sequencing. gRNAs prioritized for further analysis were selected by the fold depletion of abundance in sorafenib-treated sample compared with that in untreated sample.

### CRISPR gRNA generation and lentiviral transduction

For construction of an sgRNA-expressing vector, DNA oligonucleotides (Invitrogen) were annealed and ligated into BsmBI-digested LentiCRISPRv2 plasmid. Target sgRNA oligonucleotide sequences are listed as follows:

sgERK2-1F: 5’-CACCGCAACCTCTCGTACATCGGCG-3’;

sgERK2-1R: 5’-AAACCGCCGATGTACGAGAGGTTGC-3’;

sgERK2-2F: 5’-CACCGCGCGGGCAGGTGTTCGACGT-3’;

sgERK2-2R: 5’-AAACACGTCGAACACCTGCCCGCGC-3’.

Production of lentivirus was performed as previously described (Prahallad et al, 2012). Then, HCC cells were infected with lentiviral supernatants using 8 μg/ml Polybrene. After 24h of incubation, the supernatant was replaced by medium containing 2 μg/ml Puromycin. After 48h, selection of viral transduced cell lines was completed.

### Long-term cell proliferation assays

Cells were seeded into 6-well plates (1.5-3 × 10^4^ cells per well) and cultured in the presence of drugs as indicated. Within each cell line, cells cultured at different conditions were fixed with 4% paraformaldehyde (in PBS) at the same time. Afterwards, cells were stained with 0.1% crystal violet (in water).

### Protein lysate preparation and western blots

Cells were washed with PBS and lysed with RIPA buffer supplemented with Complete Protease Inhibitor (Roche) and Phosphatase Inhibitor Cocktails II and III (Sigma). All lysates were freshly prepared and processed with Novex NuPAGE Gel Electrophoresis Systems (Invitrogen).

### IncuCyte^®^ cell proliferation assay and apoptosis assays

Indicated cells were cultured and seeded into 96-well plates at a density of 1000-1500 cells per well. 24 hours later, drugs were added at indicated concentrations. Cells were imaged every 4 hours in IncuCyte ZOOM (Essen Bioscience). Phase-contrast images were collected and analyzed to detect cell proliferation based on cell confluence. For cell apoptosis, IncuCyte^®^ Caspase-3/7 green apoptosis assay reagent was also added to culture medium and cell apoptosis were analyzed based on green fluorescent staining of apoptotic cells.

### Immunohistochemical staining

HCC specimens were obtained from 78 patients who underwent curative surgery in Eastern Hepatobiliary Hospital of the Second Military Medical University in Shanghai, China. Patients were not subjected to any preoperative anti-cancer treatment. Ethical approval was obtained from the Eastern Hepatobiliary Hospital Research Ethics Committee, and written informed consent was obtained from each patient. Immunohistochemical staining for p-ERK was done as previously described (Wang et al, 2012).

### Xenografts

All animals were manipulated according to protocols approved by the Shanghai Medical Experimental Animal Care Commission. Huh7 and HepG2 cells (5 × 10^6^ cells per mouse) were injected subcutaneously into the right posterior flanks of 6-week-old BALB/c nude mice (6 mice per group). Tumor volume based on caliper measurements was calculated by the modified ellipsoidal formula: tumour volume = ½ length × width. When tumors reached a volume of approximately 100 mm^3^, mice were randomly assigned to treatment with vehicle, sorafenib (30 mg/kg, daily gavage), selumetinib (20 mg/kg, daily gavage), or a drug combination in which each compound was administered at the same dose and scheduled as single agents. Tumor tissues were fixed, embedded, and sliced into 5 μm thick sections. Immunostaining of p-ERK, caspase 3 and PCNA was carried out as described previously (Wang et al, 2012).

### Statistics

The data are presented as mean with SEM or SD. All graphs were plotted and analyzed with GraphPad Prism 5 Software. P-values < 0.05 were considered statistically significant.

## Acknowledgements

We thank Xiangjun Kong for the kind gift of sgERK2 vectors. This work was supported by grants from the Cancer Genomic Center Netherlands, the Dutch Cancer Society (KWF), the National Key Basic Research Program of China (973 Program: 2015CB553905), the National Natural Science Foundation of China (81672933, 81702838), Shanghai Jiao Tong University School of Medicine (YG2014MS44 and PYXJS16-004), and Shanghai Municipal Commission of Health and Family Planning (2017YQ064 and 201640007).

## Author Contributions

Cun Wang and Haojie Jin were responsible for data acquisition and analysis and drafting the article; Dongmei Gao performed the *in vivo* experiments; Liqin Wang and Roderick L. Beijersbergen provided advice for CRISPR screen; Bastiaan Evers was responsible for data analysis of CRISPR screen. Zheng Xue provided advice for the design of the study. Guangzhi Jin provided clinical samples; and René Bernards and Wenxin Qin were responsible for the conception, design, and supervision of the study. All contributing authors gave final approval for the version to be published.

